# Progressive change in expression of killer-like receptorsr and GPR56 expression defines the cytokine production potential of human CD4^+^ T memory cells

**DOI:** 10.1101/191007

**Authors:** Kim-Long Truong, Stephan Schlickeiser, Katrin Vogt, David Boës, Julia Schumann, Christine Appelt, Mathias Streitz, Gerald Grütz, Christian Meisel, Stefan Tomiuk, Julia K. Polansky, Andreas Pascher, Igor Sauer, Undine Gerlach, Birgit Sawitzki

## Abstract

Memory T cells mount an accelerated response upon re-challenge but are heterogeneous in phenotype and function. Traditionally memory T cells were classified into central memory, effector memory and terminally differentiated effector memory (T_EMRA_) cells based on expression of CCR7 and CD45RA. Functional heterogeneity even within these subsets demonstrated the need for more suitable markers. We applied bulk and single gene expression profiling of human CD4^+^ memory T cells and identified surface markers, KLRB1, KLRG1, GPR56 and KLRF1, allowing classification into “low”, “high” or “exhausted” cytokine producers. In contrast to common understanding KLRG1 expression was not associated with exhaustion and highest production of multiple cytokines was observed in KLRB1^+^KLRG1^+^GPR56^+^ T cells. Only additional KLRF1 expression was associated with a decline in cytokine production. The superiority of KLRF1 to define exhausted cytokine producers compared to classical T_EMRA_ identification was best exemplified for intrahepatic T cells in patients with inflammatory liver diseases.

## INTRODUCTION

CD4^+^ helper T cells coordinate the immune response against invading pathogens and malignancies (1-5). However, they also play a pathological role in the development of various inflammatory and/or autoimmune disorders (6, 7).

In association to their differentiation state, CD4^+^ T cell populations vary in their migratory behaviour, cytokine production potential, proliferation capacity and effector function and thus, in their overall potential to provide immediate protection upon antigen encounter (8, 9). Naïve T cells (T_N_) develop in the thymus and migrate into secondary lymphoid organs where, upon primary antigen encounter, they provide delayed effector functions and differentiate into memory T cells (10, 11). In contrast, memory T cells show an accelerated and intensified response to antigen re-encounter, which results in rapid antigen clearance. Recent findings have shown that the functional repertoire of memory T cells is manifold and subpopulations varying in their location, protection capacity and longevity have been described (10, 11).

Due to this complexity, researchers have urged to identify phenotypic properties which help to distinguish different memory T cell subpopulations (12, 13). Based on the expression patterns of lymph node homing receptors (CD62L or CCR7) and CD45 splice variants, CD8^+^ and CD4^+^ T cells were classified into CD45RA^+^CCR7^+^ naïve (T_N_), CD45RA^-^CCR7^+^ central memory (T_CM_), CD45RA^-^ CCR7^-^ effector memory (T_EM_) and CD45RA+CCR7^-^ terminally differentiated effector memory (T_EMRA_) cells (14-16). While the developmental relationship between these memory subsets is still a matter of debate, our own recent epigenomic and transcriptomic characterizations of human circulating CD4^+^ T cell subsets support a linear differentiation model in the order of T_N^-^_T_CM^-^_T_EM^-^_ Temra cells (17).

T_CM_ cells circulate between blood and lymphoid compartments like T_N_ cells and have a high proliferation potential and self-renewal capacity. Therefore, they form a long-lasting reservoir of memory T cells but display only limited protection potential as their effector cytokine production after re-activation is limited. Both CCR7^-^ T_EM_ and T_EMRA_ subsets are excluded from lymphatic organs and migrate via the blood to peripheral tissues (15, 16) where they participate in immediate protection from re-occurring infections (9, 18). This is mediated by a heterogeneous, but generally high multi-functional cytokine production potential of T_EM_ cells (9). However, contrasting findings have been published for T_EMRA_ cells, reporting either high or low cytokine production potential. The latter one being attributed to an “exhausted” phenotype of the terminally differentiated T_EMRA_ population (9, 19-21). Indeed, T_EMRA_ cells have characteristics of end-stage differentiation and thus a very low proliferative potential (19, 21, 22), they do acquire KLRG1 and CD57 but loose CD28 and CD27 expression (21). However, acquisition of CD57 and loss of CD28 or CD27 expression is not exclusive for T_EMRA_ cells but also observed for some T_EM_ cells (22, 23). These results demonstrate, that T_EM_ and T_EMRA_ populations defined by the CD45RA/CCR7^-^ classification seem to represent heterogeneous pools of cells which are functionally and phenotypically not clearly defined and distinguishable. This is particularly true for CD4^+^ memory T cells.

Due to their high pro-inflammatory cytokine production potential, CD4^+^ memory T cells are key promoters of chronic inflammation when the physiological regulatory circuits fail. Therefore, it is not surprising that increased proportions and absolute numbers of T_EM_ and also T_EMRA_ cells have been observed in patients suffering from chronic inflammatory diseases (24, 25).

However, due to the high cellular heterogeneity and possible partial functional overlap of these two populations, it seems likely that a functionally similar T cell subset with high cytokine secretion properties might be driving the chronic inflammation irrespective of their CD45RA/CCR7 phenotype. Therefore, in this study, we aimed at characterizing the true functional heterogeneity of human CD4^+^ T cell subsets at the single cell level including the identification of reliable surface markers correlating with their cytokine production properties, contributing to inflammatory diseases.

We addressed these questions by performing gene expression profiling of purified human CD4^+^ T_N_, T_CM_, T_EM_ and T_EMRA_ cells from peripheral blood of first healthy individuals followed by single cell expression studies and identified different combinations of the surface markers KLRB1, KLRG1, GPR56 and KLRF1 suitable to describe the development of human CD4^+^ memory T cells with varying cytokine production potential. Co-expression of KLRB1 with either KLRG1 or GPR56 or co-expression of all three markers was associated with high TNF-α/IFN-γ co-expression potential while additional acquisition of KLRF1 expression during terminal differentiation resulted in a reduction of the cytokine production capacity. This novel classification allowed a more precise definition of functional states of “high” and “exhausted” cytokine producers as compared to the classical T_EM_ or T_EMRA_ gating, respectively. Importantly, this could be confirmed for blood and especially intrahepatic CD4^+^ T cells from patients with inflammatory liver diseases. With these data we introduce a novel surface marker classification scheme which more precisely defines functionally distinct memory T cell subsets as compared to the CD45RA/CCR7-based categorisation. These results highlight that human memory T cell populations especially within inflamed tissues are heterogeneous and require detailed characterizations on the single cell level to identify disease-driving subsets as targets for novel therapeutic approaches.

## RESULTS

### Identification of overlapping gene signatures enriched in human CD4^+^ T_EM_ and T_EMRA_ cells

In order to get more insights into common versus different phenotypic and functional properties of CD4^+^ T_EM_ and particularly T_EMRA_ cells we performed a comparative gene expression profiling of sorted CD4^+^ T_N_, T_CM_, T_EM_ and T_EMRA_ cells from peripheral blood of healthy individuals. We focussed on genes encoding cell surface proteins in order to identify phenotypic markers associated with distinct functional properties (e.g. cytokine production potential) of different T cell subsets. Figure 1a shows a heatmap of genes identified through an intersection analysis between T_EMRA_ vs. T_N_, T_EMRA_ vs. T_CM_, T_EMRA_ vs. T_EM_ and T_EMRA_ + T_EM_ vs. T_N_ + T_CM_ cells. Thereby, we identified 38 genes that were highest expressed in T_EMRA_ cells but also significantly increased in both CD4^+^ T_EM_ as well as T_EMRA_ cells compared to CD4^+^ T_N_ and T_CM_ cells (figure 1a and supplementary table 1).

**Figure 1:**
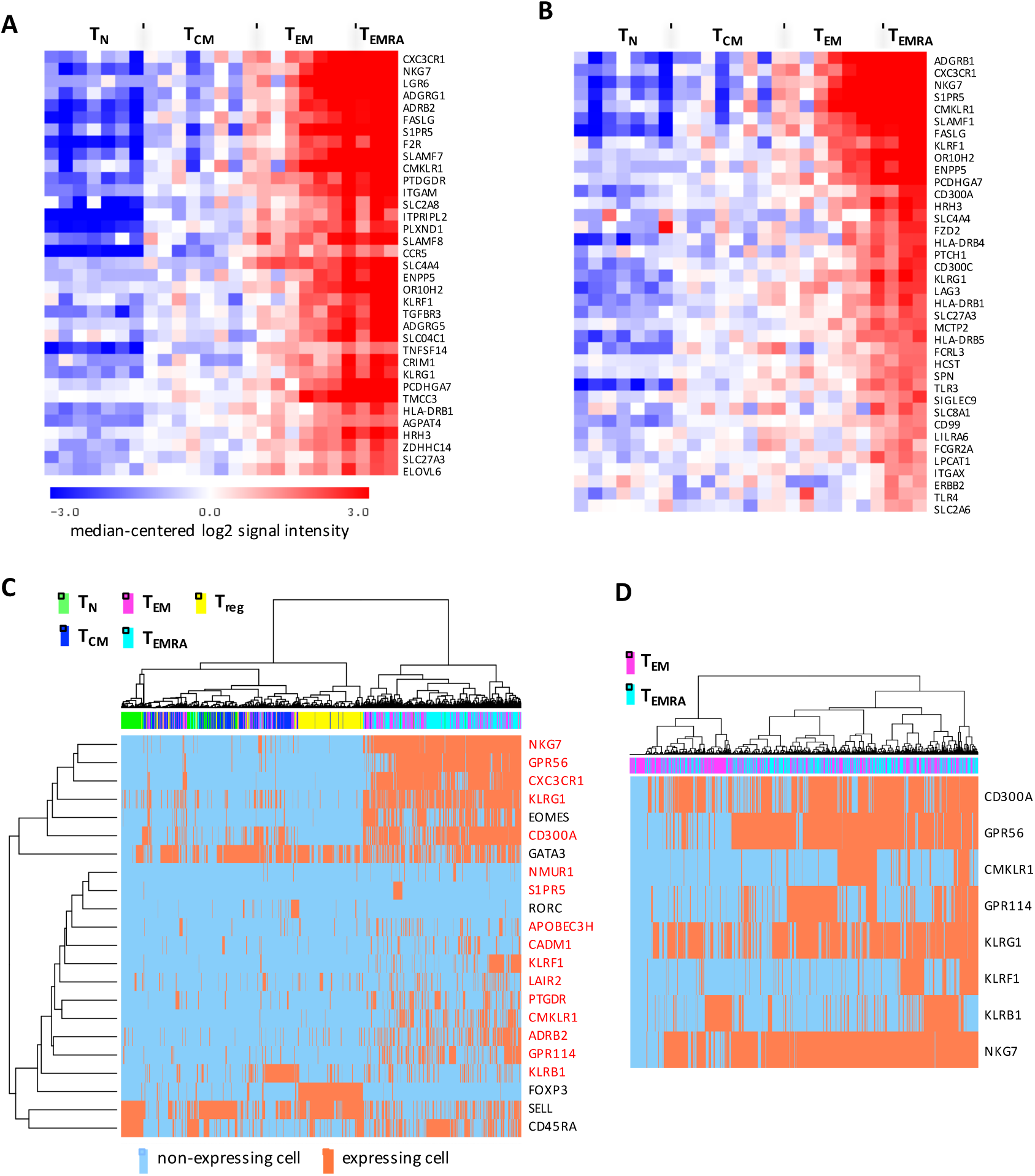
Specific mRNA expression profile and heterogeneity of CD4^+^CD45RA^-^CCR7^-^T_EM_ and CD4^+^CD45RA^+^CCR7” T_EM_ra cells. A) Heatmap of genes identified through an intersection analysis between T_EMRA_ vs. T_N_, T_EMRA_ vs. T_CM_, T_EMRA_ vs. T_EM_ and (B) heatmap of genes identified through an intersection analysis between T_EMRA^+^_ T_EM_ vs. T_N_ + T_CM_ cells comparisons of RNA microarray results of sorted CD4^+^ T_N_, T_CM_, T_EM_ and T_EMRA_ cell samples from peripheral blood (n=3-8 pools each consisting of 1-3 sorted samples). (C) Single cell profiling of T_EM_ & T_EMRA_ specific gene expression, unsupervised cluster analysis of 16 gene candidates, shown in red, and additionally selected genes in single blood CD4^+^ T_N_, T_CM_, T_EM_ T_EMRA_ cells separated from n=4 different healthy donors (total of 209 T_N_, 260 T_CM_, 396 T_EM_ 248 T_EMRA_ cells excluding debris and doublets). (D) Unsupervised cluster analysis of single cell gene expression results from identified NK cell-associated markers in blood CD4^+^ T_EM_ and T_EMRA_ cells of healthy individuals revealing homogeneous (e.g. NKG7) or heterogeneous (e.g. KLRF1) expression pattern.

Among the top-ranked genes up-regulated in both CD4^+^ T_EM_ and T_EMRA_ cells were genes previously shown to be highly expressed in more differentiated T cells such as *FASLG* or *LAG3*, genes encoding various toll-like receptors or *HLA-DR beta chains* but especially genes associated with NK cells such as members of the killer-like receptor family, *KLRG1* and *KLRF1*, or *NKG7*, *CD300A* and *CD300C.* In addition, genes encoding proteins regulating cell migration and adhesion such as *S1PR5*, *CXC3CR1* and *ADGRG1* (also known as *GPR56*) were highly up-regulated. The microarray analysis also identified genes, whose expression was specifically increased in T_EM_ cells such as genes encoding c-Kit or the killer-like receptor *KLRB1* (see also supplementary table 1 – last 6 genes). Taken together, we could identify a set of genes encoding surface markers, which were significantly higher expressed in more differentiated human CD4^+^ T_EM_ and T_EMRA_ cells.

### Heterogeneity in candidate gene expression within T_EMRA_ and T_EM_ cells

We used the identified CD4^+^ T_EM^-^_ and T_EMRA^-^_ specific genes to investigate, whether their expression was homogeneous or could be attributed to certain cell subsets within CD4^+^ T_EM^-^_ and T_EMRA_ cells. For this, we applied single cell separation combined with gene candidate-specific qRT-PCR. Separated CD4^+^ T_N_, T_CM_, T_EM_, T_EMRA_ cells as well as CD4_+_CD25^high^CD127^low^ regulatory T cells (Treg) from four healthy individuals were analysed allowing comparative gene expression analysis of a total of 209, 260, 396, 248 and 276 single cells, respectively. In addition, to the expression of the identified gene candidates, shown in red, we also analysed expression of *TBX21*, *GATA3*, *EOMES*, *RORC*, *FOXP3*, *SELL* and *CD45RA*. A complete list of all analysed genes is provided in supplementary table 2. We performed an unsupervised cluster analysis of all genes giving a signal in at least 10 cells and showing no cross-reactivity with genomic DNA (figure 1b).

Nearly all of the T_EMRA_ and the majority of the T_EM_ cells clustered separately from the other T cell subsets, which was in part due to the inclusion of lineage specific genes such as *EOMES* (figure 1b). Also, as expected, Treg cells clustered separately in a highly homogeneous cluster, underlining the clear separation of this immunosuppressive subset from all other proinflammatory subsets. A high proportion of T_EM_ but also T_EMRA_ cells transcribed *EOMES* and also *GATA3*, whereas *RORC* was only expressed by a minor fraction of T_CM_, T_EM_ and Treg cells but not by T_EMRA_ cells (suppl. table 3).

Most importantly, we could validate the selective expression pattern of most of the gene candidates at the single cell level. However, we also observed differences compared to the bulk analysis. In the bulk analysis (microarray) *KLRB1* transcription was three‐ and two-fold higher in T_EM_ cells versus T_N_ and T_EMRA_ cells, respectively (supplementary table 1). In contrast, upon single cell profiling the proportion of *KLRB1* transcribing T_CM_ cells appeared to be even higher than that of T_EM_ cells (supplementary table 3). Furthermore, only a few CD4^+^ T_EM^-^_ & T_EMRA_-specific genes such as *NKG7*, *GPR56* or *KLRG1* were expressed by nearly all T_EM_ and T_EMRA_ cells (figure 1b, c). In contrast, while the majority of genes showed a more heterogeneous expression pattern with only a fraction (e.g. *KLRF1, CMKLR1, ADRB2)* or sometimes even a minority of CD4^+^ T_EM^-^_ or T_EMRA_ cells transcribing it (e.g. *S1PR5*, *CADM1*). The variability in NK cell-associated marker expression (figure 1c) was especially apparent. As we were especially interested in understanding heterogeneity most likely associated with variations in functionality and differentiation status of the T cells, we investigated those further in more detail at protein level. We concentrated on KLRB1, KLRG1, GPR56 and KLRF1 as we achieved reproducible high-resolution antibody staining in flow cytometry (figure 2a).

**Figure 2:**
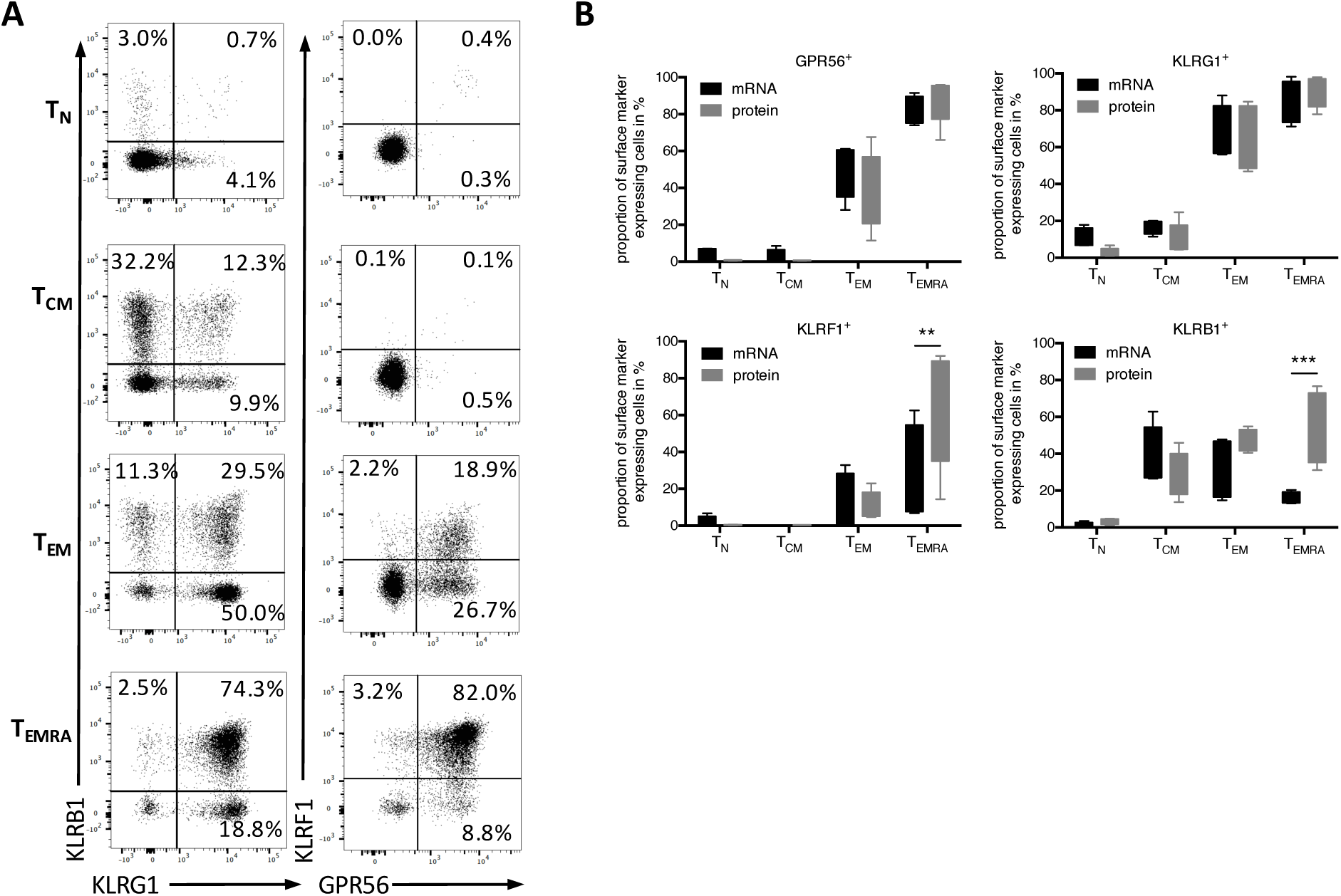
Heterogeneous surface expression pattern of killer-like receptors and GPR56 in human circulating CD4^+^ T cell subsets. (A) Validation at protein level - flow cytometry results as exemplary dot plots of KLRB1, KLRG1, KLRF1 and GPR56 expression in CD4^+^ T_N_, T_CM_, T_EM_ and T_EMRA_ cells from blood of healthy donors. (B) Boxplots comparing frequency of KLRB1, KLRG1, KLRF1 and GPR56 positive cells within CD4^+^ T_N_, T_CM_, T_EM_ and T_EMRA_ cells between single cell qRT-PCR (mRNA, n=4 separations resulting in a total of 209 T_N_, 260 T_CM_, 396 T_EM_ and 248 T_EMRA_ cells) and flow cytometry (protein, n=5) analysis. **p<0.01, ***p<0.001 Two-way ANNOVA & Sidak’s multiple comparison test (A) Exemplary dot plots revealing association of surface marker expression with cytokine expression potential of CD4^+^ T cells from peripheral blood of healthy individuals. (B) Representative t-SNE plots showing surface marker and cytokine expression pattern of pre-gated non Treg CD4^+^T cells (excluding CD25^hlgh^CD127^l0W^ cells). (C) Wanderlust analysis based on the trajectory of CD45RA and CCR7. Relative median surface markerand intracellular cytokine expression within CD4^+^T cells from blood of four healthy individuals upon short-term PMA/lono stimulation is shown as described within materials and methods.

We detected a successive increase in expression of all markers starting from T_N_ to T_EMRA_ cells with KLRB1 being already up-regulated at an early memory stage (T_CM_ cells) but expression of the other three marker occurring later during memory / effector cell development. GPR56 and especially KLRF1 up-regulation occurred at late differentiation stages. CD4^+^ T_EMRA_ cells contained the highest frequencies of KLRG1 and GPR56 as well as KLRF1 and KLRB1 double expressing cells, whereas CD4^+^ T_EM_ cells were characterised by double but also KLRB1, KLRG1 and GPR56 single expressing cells (figure 2a). Thus, we could validate the selective expression pattern of killer-like receptors and GPR56 in T_EM_ and T_EMRA_ cells. However, in concordance with the single cell mRNA expression analysis, the expression patterns where still heterogeneous and not all of the T_EM_ and T_EMRA_ cells stained positive for these four markers.

When comparing the single cell expression data on mRNA and protein level, we observed nearly identical frequencies for GPR56 and KLRG1 expression (figure 2b). In contrast, proportion of KLRF1 and especially KLRB1 positive cells varied significantly for T_EMRA_ cells pointing to fluctuations in gene transcription.

### Association of killer-like receptor and GPR56 expression with cytokine production potential of peripheral CD4^+^ T cells

Next, we tested a potential correlation between the cytokine producing potential of CD4^+^ T cells and the expression of KLRB1, KLRG1, GPR56 and KLRF1. For this, PBMCs of healthy individuals were isolated and a short-term stimulation using PMA/Ionomycin was performed, followed by surface marker and intracellular cytokine staining for acquisition by multi-parametric flow cytometry. This analysis revealed the general trend that marker positive cells display an increased potential to produce TNF-α and IFN-γ, with KLRG1 identifying the highest frequency of TNF-α & IFN-γ double producers (figure 3a).

**Figure 3:**
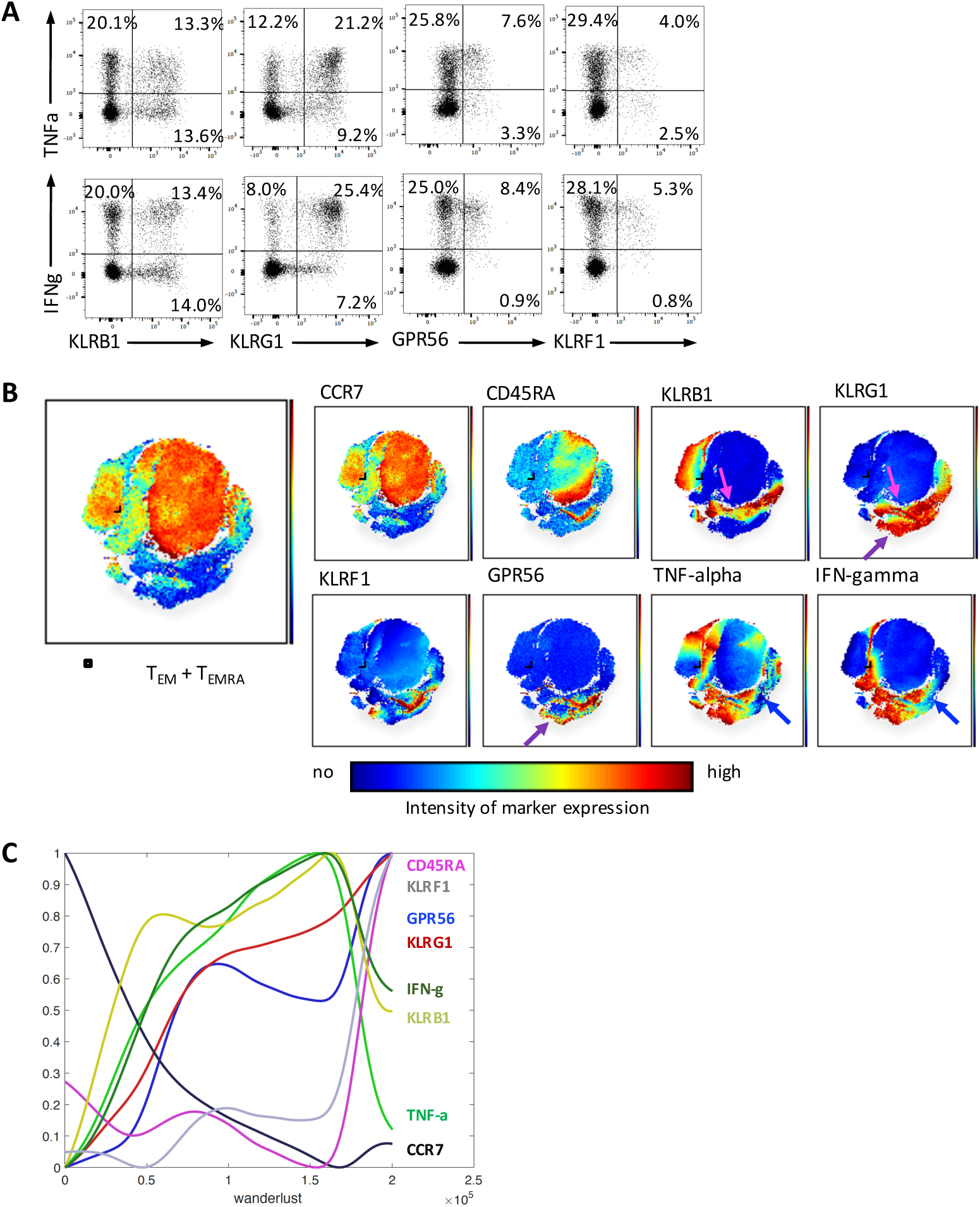
Progressive expression of killer-like receptors (KLR) and GPR56 expression and association with cytokine production potential during CD4^+^ memory T cell development. (A) Exemplary dot plots revealing association of surface marker expression with cytokine expression potential of CD4^+^ T cells from peripheral blood of healthy individuals. (B) Representative t-SNE plots showing surface marker and cytokine expression pattern of pre-gated non Treg CD4^+^T cells (excluding CD25^high^CD127^low^ cells). (C) Wanderlust analysis based on the trajectory of CD45RA and CCR7. Relativemedian surface marker and intracellular cytokine expression within CD4^+^ T cells from blood of four healthyindividuals upon short-term PMA/Iono stimulation is shown as described within materials and methods.

In order to better visualize co-expression patterns of the surface markers associating with cytokine producing potential, we created t-SNE maps arranging all conventional CD4^+^ T cells (excluding CD25^high^CD127^low^ Tregs) according to their similarity in surface marker and cytokine expression (figure 3b). We highlighted the area of CD4^+^ T_EM_ & T_EMRA_ cells within the plots (encircled black area) judging from their CD45RA and CCR7 expression pattern. As can be seen, cytokine production is common but clearly heterogeneous within the T_EM_/T_EMRA_ area, with certain subtypes being completely devoid of cytokine expression potential (blue arrows). Surprisingly, most cells in this cytokine-low area express all four surface markers with KLRF1 displaying an almost exclusive expression in this subset. Furthermore, areas of high cytokine production (TNF-α^+^ & IFN-γ^+^) contain cells which either co-express KLRB1 and KLRG1 (pink arrows) or KLRG1 and GPR56 (purple arrows). These results from visual inspection of the t-SNE maps indicated that different combinations of surface markers are characteristic for different functional states. As the acquisition or loss of cytokine expression potential is generally linked to the differentiation state of T cells, we wanted to analyse how the expression of our surface markers correlates to the differentiation pathway of memory T cells according to the CD45RA/CCR7-based classification. For this, we applied the recently described wanderlust algorithm to construct a trajectory of CD4^+^ T cell differentiation based on the classical surface marker CD45RA and CCR7 and our identified surface marker set (26). Using CD45RA and CCR7 expression we defined CD45RA^+^CCR7^+^ (T_N_) cells as the “initiator” and CD45RA^+^CCR7^-^ (T_EMRA_) cells as the “terminal” cells. We then examined the relative expression pattern of our identified marker but also intracellular TNF-α and IFN-γ along the developmental trajectory by plotting them against the wanderlust axis (figure 3c). According to this analysis, KLRB1 expression was the first marker to be acquired during CD4^+^ memory T cell differentiation, a result which is nicely confirmed by our bulk and single cell-based gene expression analyses (figures 1 and 2). Subsequently, cells started to up-regulate KLRG1 followed by a nearly induction of GPR56. KLRF1 expression was only acquired at a late stage during memory T cell differentiation. Interestingly, simultaneously to the up-regulation of KLRB1 T cells obtained the potential to produce TNF-α and with a slight delay also IFN-γ. Whereas KLRB1 and KLRG1 showed a nearly constant increase in expression during differentiation, GPR56 and KLRF1 expression followed a two-phase pattern. Late stage differentiated CD45RA re-expressing CD4^+^ T cells acquired very high KLRG1, GPR56 and KLRF1 expression but a reduction in KLRB1 expression concurrent with a decline in TNF-α and IFN-γ production.

### Combinations of different KLRs and GPR56 allow refined description of memory CD4^+^ T cell states and define magnitude of cytokine production potential

From our wanderlust analysis we concluded a progressive acquisition of our surface markers during memory T cell differentiation in the following order: KLRB1, KLRG1, GP56, KLRF1. Based on this, we analysed whether the T cell subsets defined by this new scheme would indeed recapitulate or even refine the known correlation to cytokine expression potential from "low" (classically gated T_N_ cells) to "high" (T_EM_ cells) and finally to "exhausted" (T_EMRA_ cells, Fig. 4a, top row).

**Figure 4:**
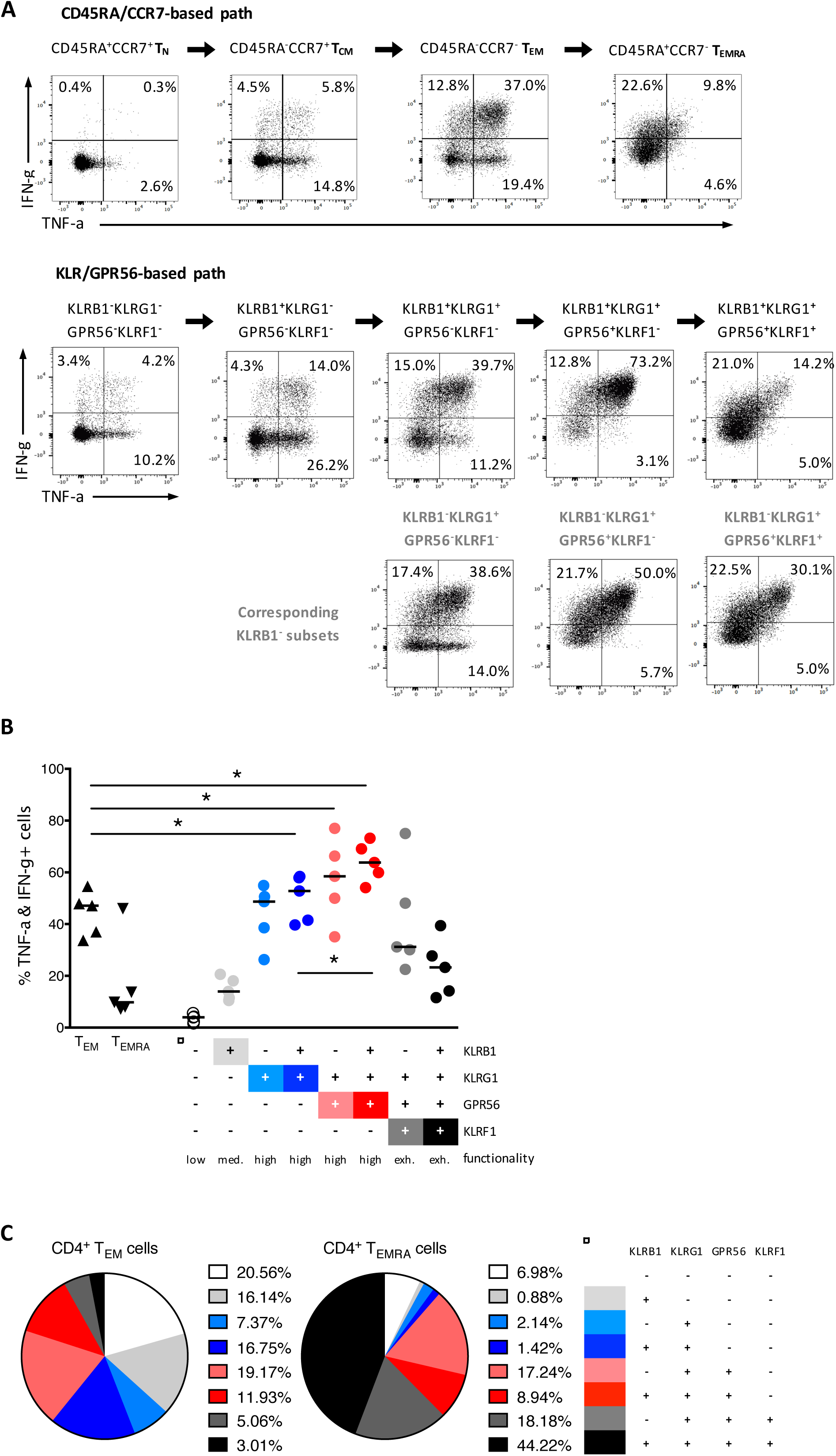
Combinational expression analysis of killer-like receptors and GPR56 defines magnitude of cytokine production and recapitulates human CD4^+^ memory T cell development. (A) Exemplary dot plots revealing changes in TNF-α and IFN-γ production potential upon short-term PMA/lono stimulation within conventionally gated (CD45RA/CCR7-based path) or KLR/GPR56-based gated CD4^+^ memory T cells. The plots are ordered according to the anticipated developmental pathway. (B) Comparative analysis of TNF-α/IFN-γ co-producing cell frequencies and functionality of conventionally gated T_EM_ and T_EMRA_ cells and newly defined subsets according to KLRB1, KLRG1, GPR56 and KLRF1 expression pattern upon short-term PMA/lono stimulation (n=5). (C) KLR/GPR56-based subset composition within classically gated CD4^+^ T_EM_ and T_EMRA_ cells (n=5). *p<0.05 (Wilcoxon matched pairs signed-rank test).

To this end we defined the following subsets within total CD4^+^ T cells: 1) no marker expression = KLRB1^-^KLRG1^-^GPR56^-^KLRF1^-^, 2) KLRB1^+^KLRG1^-^GPR56^-^KLRF1^-^, 3) KLRB1^+^KLRG1+GPR56^-^KLRF1^-^, 4) KLRB1^+^KLRG1^+^GPR56^+^KLRF1^-^ and 4) KLRB1^+^KLRG1^+^GPR56^+^KLRF1^+^ and analysed their IFN-γ and TNF-α production potential to compared it to that of classically gated T_N_, T_CM_, T_EM_ and T_EMRA_ cells (figure 4a). Indeed, the KLRs/GPR56-based subset definition allowed identification of CD4^+^ memory T cells with a continuous gain in cytokine production potential.

Interestingly, after the primary acquisition of the initial marker KLRB1, the expression of this marker seemed to contribute little to the cytokine expression potential as KLRB1^+^KLRG1^+^GPR56^-^ KLRF1^-^ and KLRB1^-^KLRG1+GPR56^-^KLRF1^-^ as well as KLRB1^+^KLRG1^+^GPR56^+^KLRF1^-^ and KLRB1^-^ KLRG1^+^GPR56^+^KLRF1^-^subsets displayed only minor differences in cytokine production potential. In fact, the KLRB1^-^ subsets (depicted in lighter colours in figure 4a) recapitulated the progressive acquisition of cytokine expression potential with memory T cell differentiation and terminal exhaustion with acquisition of the KLRF1 marker (figure 4a). Based on these results, we conclude that the combinatory expression profile of KLRB1, KLRG1, GPR56 and KLRF1 allows a refined classification of memory T cell subsets along their differentiation line and correlating to their functional state judged from their cytokine expression potentials “low”, “medium”, “high” and “exhausted” (figure 4b). This refinement now facilitates the definition of the most potent cytokine producing subsets KLRB1^+^KLRG1+GPR56^-^KLRF1^-^, KLRB1^-^KLRG1^+^GPR56^+^KLRF1^-^ and especially KLRB1^+^KLRG1^+^GPR56^+^KLRF1^-^, which containing significantly more TNF-α/IFN-γ co-producing cells as compared to traditionally gated T_EM_ cells (figure 4b).

Having revealed that T_EM_ cells contain less TNF-α & IFN-γ co-producing cells as compared to the most potent subsets with the new classification, we wondered whether indeed T_EM_ cells are composed of different subsets according to our KLRF/GPR56-based definition. Indeed, although the “high” cytokine producing subsets made up the majority of T_EM_ cells, populations with a “low” or “exhausted” functional state were also present, which may explain the overall lower cytokine production potential in T_EM_ cells (figure 4c). Furthermore, T_EMRA_ cells were composed of mainly “exhausted” populations with some of the other subsets remaining (figure 4c), but showed in general a lower cytokine production potential (figure 4b). Thus, the refined classification of memory T cells according to the KLR/GPR56 scheme reveals functional heterogeneity in the classical T_EM_ and T_EMRA_ subsets with partially overlapping composition.

### Novel KLR/GPR56-classification reveals reduction in functionally exhausted memory T cells and increase in cytokine producers in the liver compared to the peripheral blood

In recent years it became clear that significant phenotypical and functional differences exist between circulating and intra-tissue T cells (27, 28). We therefore studied our newly defined memory T cell surface marker panel on T cells derived from human liver tissue. First, we compared the proportions of CD4^+^ T cells displaying a classical T_N_, T_CM_, T_EM_ and T_EMRA_ phenotype between blood of healthy controls (HC-B) and blood (LD-B) and liver (LD-L) from patients with inflammatory liver disease. As expected, T cell from liver samples contained the lowest proportions of T_N_ and T_CM_ cells but highest of T_EM_ cells (figure 5a). Interestingly and somewhat unexpected, the proportion of T_EMRA_ cells was in some liver samples lower than in the corresponding blood samples. Next, we performed single cell gene expression profiling of all genes listed in supplementary table 2 within sorted blood‐ and liver-derived T_CM_, T_EM_ and T_EMRA_ cells (figure 5b). Unsupervised cluster analysis of selected candidate gene marker expression resulted in a separation of two main clusters which differed in the proportion of KLRB1, GPR56, NKG7 and KLRF1 expressing cells. The left cluster in figure 5b contained the majority of KLRB1^+^ cells and was dominated by T_CM_ (blood and liver) cells with enrichment of nearly all liver T_EM_ and T_EMRA_ cells. In contrast, the majority of the blood T_EM_ and T_EMRA_ cells was contained within the right cluster, which also showed a strong enrichment for GPR56, KLRF1 and partially KLRG1 expressing cells. This indicated that there was a qualitative difference between liver and blood-derived T_EM_ and T_EMRA_ cells. Indeed, the proportion of GPR56, KLRF1 and partially KLRG1 expressing T_EM_ cells showed a tendency to be lower in liver samples, whereas the opposite was true for the proportion of KLRB1 expressing T_EM_ cells (figure 5c).

**Figure 5:**
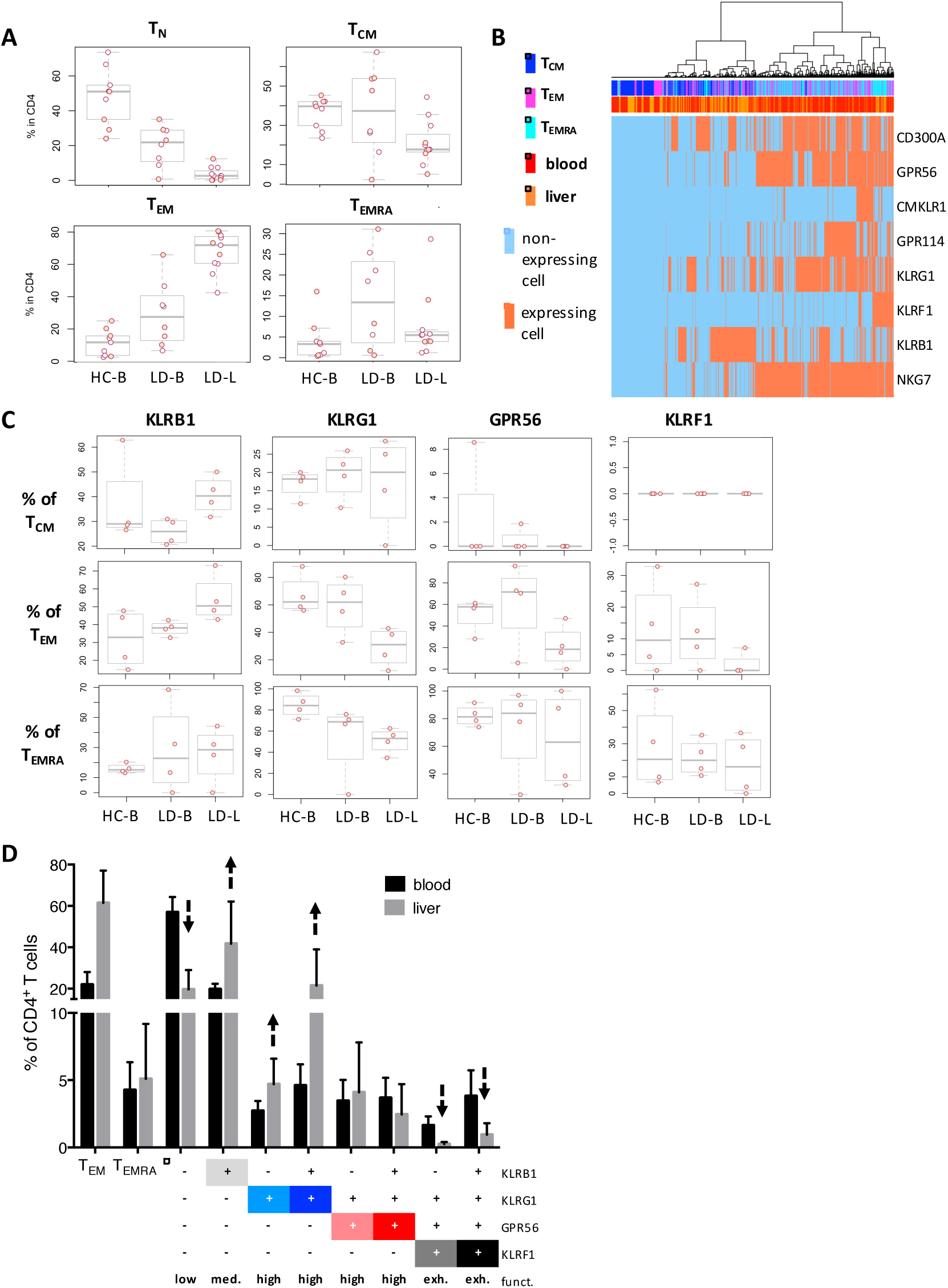
Decreased proportions of KLRF1 expressing CD4^+^ memory T cells in liver. (A) Box plots on comparative flow cytometry results of CD4^+^T_N_, T_CM_, T_EM_ and T_EMRA_ cells from blood of healthy individuals (HC-B, n=8) as well as blood (LD-B, n=8) and liver samples (LD-L, n=ll) of patients. (B) Unsupervised cluster analysis of candidate gene expression results at single cell level between blood (n=5) and liver (n=5) samples in a total of CD4^+^ T_CM_ (blood 258, liver 167), T_EM_ (blood 282, liver 201) and T_EMRA_ (blood 289, liver 238) cells. (C) Frequencies of T_CM_, T_EM_ and T_EMRA_ cells transcribing the gene markers. (D) Proportions of CD4^+^ T_EM_ and T_EMRA_ cells and newly defined subsets according to KLRB1, KLRG1, GPR56 and KLRF1 expression within CD4^+^ T cells from blood (n=5) and liver (n=5) of patients assessed by flow cytometry.

These findings led us to investigate whether the proportions of T cell subsets defined according to our novel KLR/GPR56 classification were different between blood and liver samples. Indeed, liver samples contained less of the low cytokine producing T cell subset KLRB1^-^KLRG1^-^GPR56^-^ KLRF1^-^, and increase in the high cytokine producing subsets KLRB1^-^KLRG1^+^GPR56^-^KLRF1^-^ and KLRB1^+^KLRG1^+^GPR56^-^KLRF1^-^ (figure 5d). The most striking difference was the reduction in the exhausted phenotypes KLRB1^-^KLRG1^+^GPR56^+^KLRF1^+^ and KLRB1^+^KLRG1^+^GPR56^+^KLRF1^+^. These changes in subset composition accumulate to a generally increased pro-inflammatory functionality in the liver of patients compared to the blood. In line with this, liver T_EM_ and T_EMRA_ cells showed a different subset composition based on the KLR/GPR56 classification as compared to their blood counterparts with a clear reduction of functionally exhausted KLRF1^+^ subsets in both populations (figure 6a). While the cytokine expression potential in the exhausted subsets was even lower than their blood-derived counterparts, all other subsets showed a generally increased cytokine production in the liver compared to the blood (figure 6b).

**Figure 6:**
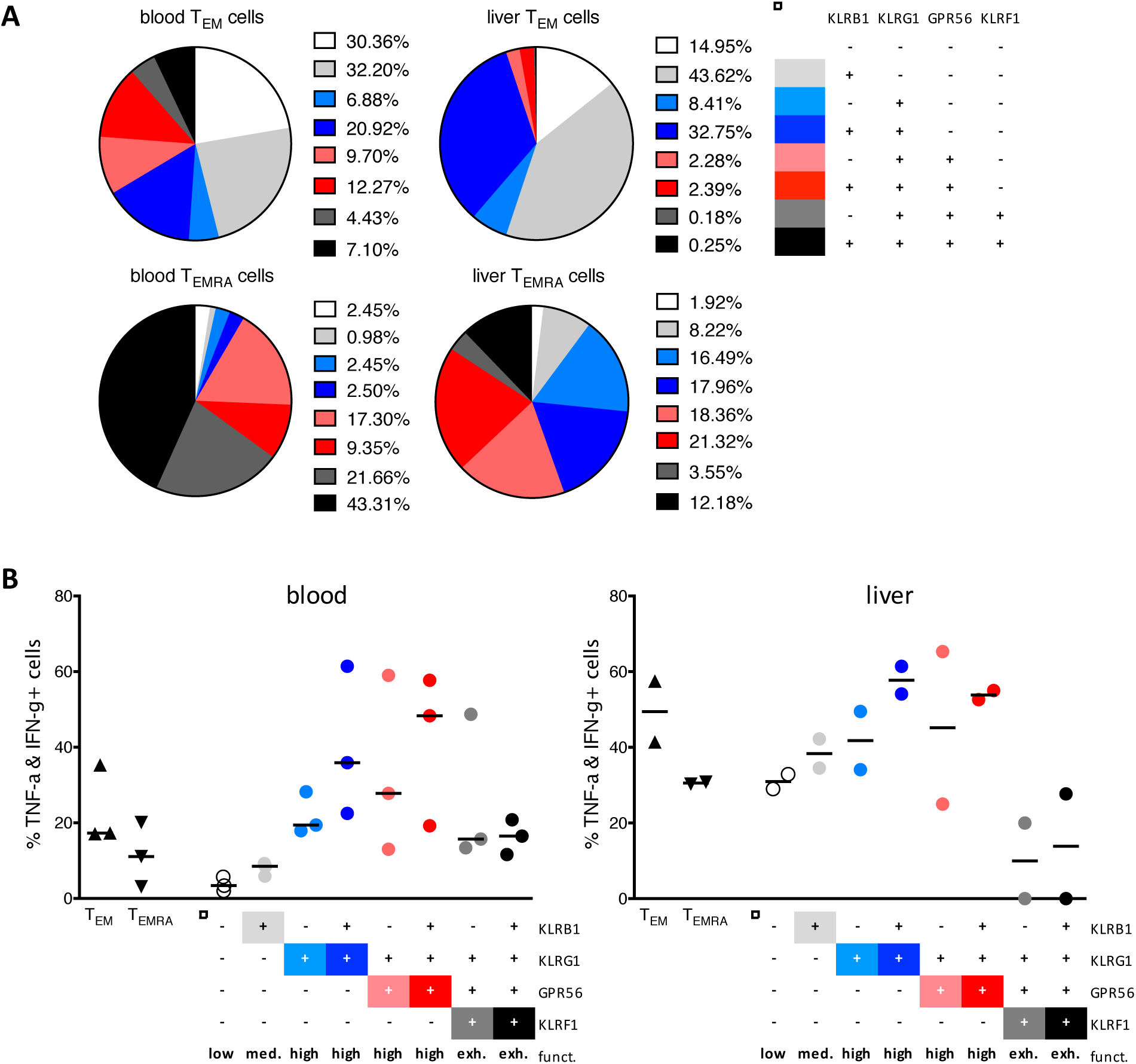
Altered subset composition and cytokine expression pattern of T_E_m and T_E_mra cells of intra-hepatic compared to peripheral CD4^+^ memory T cells. (A) Subset composition gating within CD4^+^ T_EM_ and T_EMRA_ cells of patient blood (mean, n=4) and liver samples (mean, n=4). (B) Proportions of TNF-α/IFN-γ co-expressing cells upon pre-gating of T_EM_, T_EMRA_ or newly defined subsets following PMA/Iono stimulation of PBMCs (n=3) or liver leukocytes (n=2).

Taken together, our findings introduce a novel surface marker classification scheme for CD4^+^ memory T cells, which recapitulates their differentiation pathway and more precisely indicates the cytokine production potential of each subset compared to the classical CCR7/CD45RA-based index. This might be of particular interest for the characterization of T cells from diseased tissues, in which specific functional subsets might play an essential role for the pathophysiology and maintenance of disease.

## DISCUSSION

Ever since the first categorization of CD4^+^ or CD8^+^ T cells into populations defining their differentiation status based on CD45RA and CCR7 expression discussions arose about the overall validity (9). Indeed, recent findings on functional classification of CD8^+^ memory T cells have revealed that categorisation based on CD62L or CCR7 expression and thus lymph node homing properties is not sufficient (29). Therefore, it is not surprising that it was questioned whether the defined subsets represented indeed homogeneous populations or whether individual cells differed greatly in their functional state (30-35), which for CD4^+^ T cells is mainly defined by their cytokine expression potential.

We here addressed these questions in the human system for CD4^+^ memory T cells and assessed for the first time the cellular heterogeneity on the single cell level within classically-gated CD4^+^ memory T lymphocytes from the blood as well as from liver tissue of patients. As expected, we found a pronounced heterogeneity within each subset on the overall transcriptional level, but also on the functional level assessed by single-cell cytokine secretion measurements. From these data, we developed a novel subset classification system based on the progressive acquisition of surface expression of the NK cell-associated proteins KLRB1, KLRG1, GPR56 and KLRF1. We show that this classification thoroughly mirrors the memory differentiation line and is superior in indicating the cytokine production potential of the individual subsets compared to the classical CD45RA/CCR7-based system.

Our findings on the concurrent expression of multiple KLRs and final acquisition of KLRF1 and a decline of cytokine production potential is in line with published reports on murine CD8^+^ and CD4^+^ memory T cells. Analysis of phenotypic properties of murine CD8^+^ and CD4^+^ exhausted memory T cells revealed a correlation between concurrent expression of multiple inhibitory receptors such as PD-1, LAG-3, 2B4 (CD244) and CD160 or PD-1, CTLA4, CD200 and BTLA, respectively, with decreased TNF-α/IFN-γ co-production potential (36, 37). However, investigations on human memory T cells and in particular CD4^+^ memory T cells have been not performed so far.

Also, the investigations on murine memory T cells were either limited to the characterisation of T cells with high effector function or non-functional exhausted T cells and did not allow following the complete memory T cell development. Incorporating novel technologies and analysis algorithms such as single cell gene expression profiling and Wanderlust enabled us to propose a new path of human CD4^+^ memory T cell development defining populations with “low”, “medium”, “high” and finally “exhausted” functional states. In our screen expression pattern of KLRB1, KLRG1, GPR56 and KLRF1 appeared to be most informative. Although PD-1 expression is associated with T cell exhaustion (38) and thus terminal differentiation of T cells, our initial RNA microarray analysis did not reveal a significant enrichment of PD-1 transcription within CD4^+^ T_EM_ and T_EMRA_ cells as we also observed transcription in T_CM_ cells.

Our four identified surface marker, KLRB1, KLRG1, GPR56 and KLRF1, were all first described in relation to their high expression in NK cells (18, 39-42) indicating similarities between NK cell differentiation and memory/effector T cell development.

The C-type lectin KLRB1 also known as CD161 has been shown to be expressed by CD4^+^ and CD8^+^ T cells. For CD4^+^ T cells, KLRB1 expression was mainly ascribed to IL-17 producing Th17 cells (43). However, other recent publications identified also broader KLRB1 expression across different T cell lineages expressing e.g. IL-17 or TNF-α/IFN-γ which is in agreement with our findings (44-46). For NK cells KLRB1 ligation is generally accepted to be inhibitory (44). In contrast, for T cells inhibitory as well as costimulatory roles have been proposed (44). This might explain our results as acquisition of KLRB1 expression was associated with a first significant increase in cytokine producing CD4^+^ T cells. Interestingly, we did observe differences between KLRB1 transcription and protein expression. Whereas, KLRB1 transcription was mainly limited to T_CM_ and T_EM_ cells, protein expression was observed for T_CM_, T_EM_ and T_EMRA_ cells and not down-regulated even upon acquisition of an “exhausted” cytokine production fate.

The killer cell lectin-like G1 (KLRG1) is a marker for T cell senescence as expressing cells have limited proliferative capacity (19, 21). However, KLRG1 expressing T cells are not exhausted as they display cytokine production and cytotoxic potential (47). KLRG1 expression is supposed to be limited to tissue-homing and thus T_EM_ and T_EMRA_ cells (8, 28, 48-50). Our own data showed that also a significant proportion (≈22%) of CCR7 expressing T_CM_ cells KLRG1^+^. These findings are in agreement with other published reports showing that also T_CM_ cells can express KLRG1 which was associated with increased production of effector cytokines of the expressing T_CM_ cells (35). Indeed, also our results revealed a dramatic increase of cytokine production potential as soon as the T cells acquired KLRG1 expression. This again shows the superiority of our identified KLR/GPR56-based categorisation over the traditional CD45RA/CCR7-based system.

Already in the first report describing the NK cell triggering activity of KLRF1, also known as NKp80, its expression on a subset of T cells was observed (42). In addition, it was shown that NKp80 ligation can augment CD3-stimulated degranulation and IFN-γ secretion by effector memory CD8^+^ T cells (51). This is contradictory to the here described results as KLRF1 acquisition was associated with a decline in cytokine production potential. However, our investigations were performed on CD4^+^ T cells, and for murine T cells distinct properties for CD4^+^ in comparison to CD8^+^ T cells were recently described (37). It remains to be investigated whether KLRF1 plays an inhibitory role for human CD4^+^ memory T cell activation. Nevertheless, KLRF1 expression was able to identify memory CD4^+^ T cells with reduced cytokine production potential regardless of cohort (healthy control vs. patient) or tissue type origin.

GPR56 was shown to be expressed by cytotoxic NK and T lymphocytes including CD8^+^, CD4^+^ and γd^+^ T cells (41). For NK cells an inhibitory role for GPR56 in controlling steady state activation by associating with the tetraspanin CD81 was revealed (52). Similar to KLRF1, the role of GPR56 for stimulation-dependent production of cytokines by human CD4^+^ T cells is unknown and needs to be investigated in further studies.

Our findings of successive expression of several killer-like receptors and GPR56 concurrent to first increasing and finally declining cytokine production potential is completely in line with our previous findings on linear differentiation from T_CM_, via T_EM_ and towards T_EMRA_ cells (17). There we detected an increase in global demethylation which could explain the successive expression pattern described here.

Finally, our results with increased T_EM_ but decreased or equal T_EMRA_ frequencies in liver in comparison to blood is in agreement with recent descriptions on the spatial map of human T cell compartmentalization (11). Although the authors did not investigate liver tissue, they reported also increased T_EM_ frequencies within intestinal and lung tissues in comparison to blood whereas T_EMRA_ frequencies did not vary. However, the here reported combinational expression pattern of KLRs and GPR56 challenge the analysis of overall T_EM_ and T_EMRA_ frequencies as analysis of KLRB1, KLRG1 and / or GPR56 expression versus final acquisition of KLRF1 seem to be superior to discriminate between “high” and “exhausted” cytokine producing cell subsets, which are decreased in intra-hepatic CD4^+^ T_EM_ and T_EMRA_ cells in comparison to their blood equivalents.

In summary our data reveal that identifying human CD4^+^ memory T cell populations based on the expression pattern of KLRB1, KLRG1, GPR56 and KLRF1 enables a better definition of functional states especially in peripheral tissues as compared to the classical CD45RA/CCR7-based categorisation. These findings will have enormous implications for clinical diagnostics, development of novel target-specific immune therapies as well as a better understanding of CD4^+^ memory T cell development and function. It will be interesting to see whether the here described combinational expression profile and functional subsets might aid improved prediction of disease progression in inflammatory diseases or therapeutic efficacy upon vaccination or checkpoint inhibition.

## MATERIALS AND METHODS

### Peripheral blood and liver samples

Heparinized blood and liver samples from patients who underwent liver explantation or partial liver resection as treatment for diseases of the biliary tract, including Caroli disease, gall bladder carcinoma, cholangiocellular carcinoma, Klatskin tumor and alcoholic liver cirrhosis. Median age of the patients was 67, ranging from 50 years to 79 years. Liver samples were taken from healthy, non-cancerous and non-necrotic parts of the resected liver tissue and were preserved in Hank's balanced salt solution (HBSS). Heparinized blood from age-matched healthy individuals was collected. Sample collection was performed following the Declaration of Helsinki, the European Guidelines on Good Clinical Practice, with permission from the relevant national and regional authority requirements and ethics committees (EA2/044/08 & EA1/116/13, Ethics Committee of the Charite Berlin). All samples were processed within an hour after retrieval.

### Isolation of peripheral blood mononuclear cells (PBMC)

PBMC were isolated at room temperature by density gradient centrifugation (Biocoll, Biochrom, Berlin, Germany) of heparinized blood diluted 1:2 in Phosphate-Buffered Saline (PBS) (Gibco, Thermo Fisher Scientific, Paisley, UK). Cell number was determined using a hemocytometer. Isolated PBMC were directly used for sorting, stimulation or were cryopreserved.

### Isolation of intrahepatic lymphocytes (IHL)

Liver tissue was dissected into 1 mm^3^ fragments and digested with agitation (75-80 rpm) at 37 °C for 30 minutes in a digestive solution (2 % FCS, 0.6 % bovine serum albumin, 0.05 % collagenase type IV and 0.002 % DNAse | per 1 g tissue and 10ml). Undissociated tissue was pressed through a steel sieve with a syringe plunger and dissolved in the same solution. Dissociated tissue was centrifuged at 500 × g. Tissue components were diluted in HBSS. The tissue suspension was centrifuged at 30 × g to separate and discard the formed hepatocyte-rich matrix. Still undissociated tissue was removed by filtration through 100 μm nylon mesh, leaving a cell suspension. Hepatocytes were removed using a 33 % Bicoll density gradient centrifugation. Red blood cells were lysed using water. Isolated intrahepatic lymphocytes were cryopreserved in liquid nitrogen.

### Antibody staining and T cell subset sorting

MACS-enriched CD4^+^ T cells (CD4 microbeads, human, Miltenyi Biotec) were stained in MACS-buffer (PBS with 0.5% BSA and 2 mM EDTA) at a concentration of 2 × 10^8^ cells per ml for surface expression using anti-CCR7-Alexa488 (GO43H7), anti-CD25-PE (M-A251), anti-CD45RA-PE-Cy7 (HI100), anti-CD127-APC (A019D5), anti-CD3-Alexa700 (UCHT1), anti-CD4-BV510 (OKT4), and anti-CD45R0-ECD (UCHL1, all from BioLegend). Cells were washed and stained with DAPI.

Cells were sorted using a BD FACSAria^TM^ || into the following CD4^+^ T cell subpopulations: Treg (CD25^HIGH^CD127^LOW^) and non-Treg: Tn (CD45RA^+^CCR7^+^), T_CM_ (CD45RA^-^CCR7^+^), Tem (CD45RA^-^CCR7^-^) and Temra (CD45RA^+^CCR7^-^).

### RNA microarray analysis

Total RNA from sorted T cell populations was isolated using TRIzol (Thermo Fisher Scientific, Bremen, Germany). RNA quality and integrity were determined using the Agilent RNA 6000 Nano Kit on the Agilent 2100 Bioanalyzer (Agilent Technologies). RNA was quantified by measuring A260nm on the ND-1000 Spectrophotometer (NanoDrop Technologies).

### RNA Amplification and Labeling

Sample labeling was performed as detailed in the “One-Color Microarray-Based Gene Expression Analysis protocol (version 6.6, part number G4140-90040). Briefly, 10 ng of each total RNA samples was used for the amplification and labeling step using the Agilent Low Input Quick Amp Labeling Kit (Agilent Technologies). Yields of cRNA and the dye-incorporation rate were measured with the ND-1000 Spectrophotometer (NanoDrop Technologies).

### Hybridization of Agilent Whole Mouse Genome Oligo Microarrays

The hybridization procedure was performed according to the “One-Color Microarray-Based Gene Expression Analysis protocol (version 6.6, part number G4140-90040) using the Agilent Gene Expression Hybridization Kit (Agilent Technologies). Briefly, 1.65 μg Cy3-labeled fragmented cRNA in hybridization buffer was hybridized overnight (17 hours, 65 °C) to Agilent Whole Human Genome Custom Oligo Microarrays 4×44K (AMADID 014850) using Agilent’s recommended hybridization chamber and oven. Following hybridization, the microarrays were washed once with the Agilent Gene Expression Wash Buffer 1 for 1 min at room temperature followed by a second wash with preheated Agilent Gene Expression Wash Buffer 2 (37 °C) for 1 min. The last washing step was performed with acetonitrile.

### Single-cell gene expression analysis

The C1^TM^ Single-Cell Auto Prep System (Fluidigm, South San Francisco, CA, USA) was used for single cell isolation and preamplification to prepare separate single cell cDNA in a 5-10 μm C1^TM^ Single-Cell PreAmp Integrated Fluidic Circuit (IFC) within a C1^TM^-Chip. For single-cell isolation a cell suspension of at least 660.000 cells/ml was used, which enabled at least 2000 cells to enter the C1^TM^-chip. Visualization of cell loading (empty, single, doublets or debris) was done using a light microscope. Single-cell capture rates were documented. Cell lysis, reverse transcription and preamplification was performed on the C1^TM^-chip. Afterwards cDNA of each cell was harvested for qRT-PCR preparation. CDNAs and 48 TaqMan^TM^ gene expression assays (Thermo Fisher Scientific), including RNA Spike 1 and B2M as control values, were applied to the BioMark^TM^ Gene Expression 48.48 IFC.

### PBMC and IHL stimulation and intracellular cytokine staining

5×10^6^ freshly isolated or thawed PBMC or IHL were stimulated with phorbol myristate acetate [50 ng/ml] (Sigma-Aldrich, Steinheim, Germany) and Ionomycin [1 μg/ml] (Biotrend, Cologne, Germany) for six hours (37 °C, 5 % CO_2_); Brefeldin A [10 μg/ml] (Sigma-Aldrich) was added two hours after start of stimulation. Unstimulated cells were also incubated for six hours.

Cells were washed once with DPBS and stained with Zombie UV^TM^ Fixable Viability Kit (BioLegend, San Diego, USA) for 15 min. Cells were washed once with staining buffer (DPBS with 2 %FBS and 0.1 % sodium azide (Serva, Heidelberg, Germany), subsequently treated for 5 min with Beriglobin^®^ (3μg/ml, CSL Behring, Marburg, Germany), directly surface stained with anti-CD127-APC-Alexa Fluor^®^ 750 (R34.34), anti-CD25-PC5.5 (B1.49.9) (both Beckman Coulter, Krefeld, Germany), anti-CCR7-Brilliant Violet 421^TM^ (G043H7), anti-CD45RA-Brilliant Violet 605^TM^ (HI100), anti-CD8a-PerCP (RPA-T8) (BioLegend) and anti-KLRG1-APC (REA261), anti-KLRB1-PE (191B8) and anti-KLRF1-PE-Vio770 (4A4.D10) (all from Miltenyi Biotec) for 75 min. Afterwards, cells were fixed and permeabilized (BD Cytofix/Cytoperm^TM^ Fixation and Permeabilization Solution, BD Biosciences, Heidelberg, Germany) for 20 min. After washing twice with Perm/Wash buffer (BioLegend) PBMCs were stained intracellularly with anti-TNF-α-Alexa Fluor^®^ 700 (Mab11), anti-IFN-Y-PE-Dazzle^TM^ 594 (4S.B3), anti-CD3-Brilliant Violet 510^TM^ (OKT3) and GPR56-VioBright^TM^ FITC (REA467) (Miltenyi Biotec) for 30 min. Samples were washed, and acquired on a BD LSRFortessa^TM^ (BD Biosciences). Data analysis was performed using FlowJo^TM^ software version 10.1 (FlowJo, LLC, Ashland, OR, USA).

To generate and visualize wanderlust trajectories of developmental changes in marker expression of CD4^+^ T cells we used the Matlab Cyt toolbo× (26). The algorithm was run on CD8-pre-gated PMA/Ionomycin stimulated samples. In order to apply Wanderlust to samples where all gradual differentiation states are present, FCS files were selected to have a high proportion of T_EMRA_ cells (5 healthy and 3 diseased) and density-dependent down-sampled to a total 50000 cells using the R SPADE package (53).

Utilizing the Cytobank viSNE tool, t-SNE maps were generated for CD8-pre-gated T cells (w/o CD25^high^CD127^low^ cells) of PMA/Ionomycin stimulated samples, allowing for visualization of the phenotypic and functional heterogeneity at single cell level (54, 55). CCR7, CD45RA, KLRB1, KLRF1, KLRG1, GPR56, TNF-α, IFN-γ, and CD127 were selected for both, Wanderlust and t-SNE dimension reduction.

### Statistics

To test for differences in frequencies of T cell subsets transcribing (single cell qRT-PCR) the gene marker or expressing them at protein level (flow cytometry) a two-way ANNOVA with Sidak's multiple comparison test was performed. Differences in proportions of TNF-α/IFN-γ double producers (paired samples) were tested using the non-parametric Wilcoxon matched-pairs signed rank test. P < 0.05 was considered statistically significant. Statistical analyses were calculated with GraphPad Prism 6.00 or R v3.3.1.

## AUTHOR CONTRIBUTIONS

K-L.T. and S.S. performed the experiments, analysed the data and wrote the manuscript; D.B. and J.Sch. helped with the experiments; K.V. assisted with C1 experiments; C.A. and M.St. performed or helped with the in vitro stimulation and flow cytometry experiments; G.G., C.M. and J.K.P. discussed and interpreted the data and edited the manuscript; St.T. performed the bioinformatics and statistical analysis of the microarray data; A.P., I.S. and U.G. helped in collecting patient samples and clinical data information; B.S. designed and supervised experiments, discussed and interpreted the data, contributed to the writing and editing of the manuscript. K-L.T. and S.S. performed the experiments, analysed the data and wrote the manuscript;

## ACKNOWLEDGMENTS

We thank Dr. Désirée Kunkel and Dr. Sarah Warth from BCRT Flow Cytometry Lab (BCRT-FCL) for assistance with cell sorting. This work was supported by the Deutsche Forschungsgemeinschaft (SFB650).

